# Automated Transcription Factor Cluster Analysis Toolkit (ATAT)

**DOI:** 10.64898/2026.11.23.768294

**Authors:** Samantha Fallacaro, Mustafa Mir

## Abstract

Many nuclear regulatory proteins, including transcription factors (TFs) form clusters often referred to as hubs or condensates. These clusters vary widely in their enrichment, size, and persistence, yet tools for quantifying these properties are typically developed for individual datasets. Here, we present the Automated TF Cluster Analysis Toolkit (ATAT), a Python-based toolkit for quantifying TF clustering in four-dimensional fluorescence microscopy data. ATAT combines two analytical pipelines: SCAN, which segments nuclear clusters and measures their spatial distribution, enrichment, and persistence, and LOCK, which quantifies TF enrichment and its dynamics at individual genomic loci. Using live *Drosophila* embryos, we demonstrate that ATAT distinguishes TF clusters from background fluorescence fluctuations across imaging conditions and resolves differences in TF enrichment and persistence at specific genes. ATAT provides a reproducible framework for investigating the spatial and temporal organization of transcriptional regulators in living cells.

**MOTIVATION:** Fluorescence microscopy has shown that many nuclear proteins, including transcription factors, form clusters, but how these clusters arise and their function roles in gene regulation remain poorly understood. Answering these questions requires quantitative measurements of cluster properties and dynamics, but at a nuclear and gene locus specific level. Current approaches are often tailored to specific datasets, making comparisons across experiments and proteins challenging. Furthermore, the dynamic nature of nuclear clusters makes it challenging to track their properties over time. We developed ATAT to address these challenges by combing automated 3D cluster segmentation and characterization (SCAN) with quantification of cluster enrichment and persistence at individual gene loci (LOCK). ATAT provides a reproducible way to investigate the properties of nucleus clusters in 3D live-cell imaging data and to relate it to activity at specific genes.

## INTRODUCTION

The spatiotemporal regulation of gene expression depends on the ability of sequence-specific transcription factors (TFs) and their co-factors to find and bind their target sites within the crowded nuclear environment. Many nuclear regulatory proteins form high-local concentration assemblies, termed hubs, clusters, or condensates, which in some cases are proposed to increase local binding frequency and maintain target occupancy, and in others to reflect molecular binding kinetics ^1–12^. The size and persistence of these clusters can depend on the concentration of their genomic substrates ^13,14^. Clusters formed by different proteins also differ in their stability. For example, repressive factors have been observed to form larger, less mobile, more persistent assemblies^14,15^, whereas activators tend to form more transient clusters^8,16,17^. Cluster properties are frequently altered in disease^18^. In several cancers, oncogenic mutations and fusion proteins generate clusters that are absent from normal cells or that differ in their properties, and these changes coincide with altered gene activation at the affected loci^19–26^. Being able to quantify cluster properties and link them to gene-specific transcriptional states is thus paramount to understanding their mechanisms of formation and function. While clustering may emerge through a variety of mechanisms, these do not need to be resolved before their properties can be quantified^12,27–29^. Cluster enrichment, size, and persistence can be measured directly from imaging data, and quantifying these properties in combination with experimental perturbations can be used to provide insight into their functions and mechanisms of formation.

There is currently no widely adopted standard for detecting and quantifying transcription factor clusters. Most studies rely on analysis procedures developed for individual datasets, often involving manually selected intensity thresholds, distance cutoffs, or cluster boundaries^8,17,30,31^. Other approaches use local intensity maxima or segmentation-free statistics to characterize spatial heterogeneity without defining individual clusters^10,32^. Most of these analyses rely on 2D sections, projections, or time-averaged measurements. Because these approaches measure different things, results are difficult to compare across studies, proteins, and imaging conditions, and the choice of boundary-detection method alone can change the measurements. Manual analysis is also labor-intensive and dependent on user judgment.

Following clusters over time is harder still. In dense nuclei, general-purpose trackers such as TrackMate^33,34^ and u-track^35^ struggle when clusters merge, split, or briefly dim^36^. Apparent merging or splitting events may also reflect distinct clusters moving within the resolution limit of the microscope rather than physical changes in the underlying assemblies^37^. A common alternative is to stop tracking the clusters and instead track a stable genomic marker, asking what the protein does at that site^8,10,17,38–40^. A labeled locus, such as an MS2-marked site of active transcription or a ParB-labeled gene, makes the tracked object independent of the cluster signal, so tracking no longer depends on cluster segmentation. By tracking an independently labeled genomic locus, these approaches quantify the local accumulation of a protein without requiring continuous identification of the same cluster across frames^9,37^.

To enable consistent quantification of nuclear protein clustering and its relationship to specific genomic loci, we developed ATAT, a Python-based toolkit comprising two integrated modules: SCAN and LOCK. SCAN (Segmentation and Clustering Analysis in Nuclei) uses automated nuclear masking, cluster segmentation, and 3D feature extraction to provide a comprehensive characterization of the global protein distribution. SCAN quantifies cluster densities, intensity, spatial distribution, and signal persistence. LOCK (Locus-specific Occupancy and Clustering Kinetics) was designed to quantify the relationship between TF clusters and specific genomic targets. When a genomic locus is labelled using fluorescence tagging systems (e.g. ParS/ParB, LacO/LacI arrays) or when nascent transcription is monitored using aptamer systems (e.g. MS2/MCP, PP7/PCP), LOCK calculates radial fluorescence profiles to measure the local accumulation of TFs. Crucially, by assessing overlap with SCAN-segmented clusters, LOCK provides measurements of the temporal persistence of TF accumulations at a tracked gene locus. Together, ATAT provides a unified approach for measuring protein clustering across the nucleus and at individual genes.

## RESULTS

### Overview and Input Requirements of ATAT

ATAT is a Python toolkit that quantifies TF clusters from 3D time-lapse images, both across whole nuclei (SCAN) and at a single tracked gene locus (LOCK). Its input is a 3D time-lapse of a fluorescently tagged protein, such as lattice light-sheet data (Fig. 1A). For locus analysis, LOCK also takes a second channel that marks a transcription site or genomic locus, for example MS2/MCP (Fig. 1A). The pipeline has three stages, and each consumes the output of the one before. A shared preprocessing stage deconvolves the images, segments every nucleus with Cellpose, and links nuclei across frames by nearest-neighbor tracking (Fig. 1B). SCAN then reads the tracked nuclei and deconvolved images and writes a table of clusters with their positions, volumes, enrichment, and persistence (Fig. 1B). LOCK reads the same nuclei and images plus SCAN’s cluster masks, so it cannot run on a dataset until SCAN has. LOCK then tracks the locus and measures TF intensity changes and overlap with the clusters defined by SCAN (Fig. 1C).

**Figure 1.**
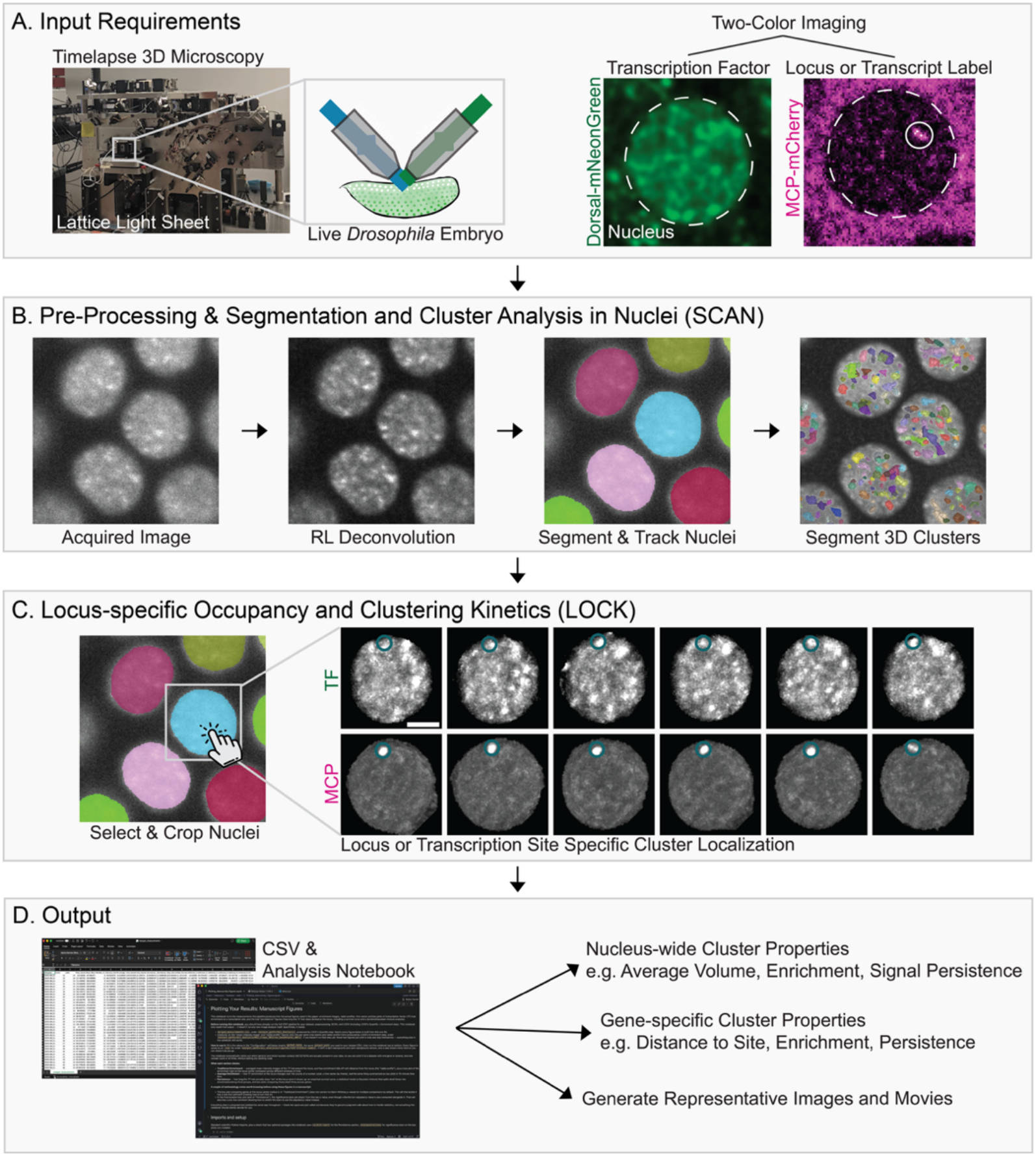
ATAT pipeline from raw imaging to locus-specific cluster measurements. (A) ATAT takes 3D timelapse fluorescence microscopy of a live embryo. As an example, we show here lattice light-sheet imaging of a *Drosophila* embryo in which a transcription factor is fluorescently tagged (Dorsal-mNeonGreen, green) and a second channel marks a transcription site or genomic locus (MCP-mCherry bound to an MS2 reporter, magenta). The solid circled spot marks that transcription site within the nucleus (dashed outline). (B) In preprocessing and SCAN (Segmentation and Clustering Analysis in Nuclei), each acquired image is deconvolved, every nucleus is segmented and tracked frame-to-frame (colored labels), and SCAN then detects and segments individual TF clusters throughout each nucleus’s volume (colored objects). (C) In LOCK (Locus-specific Occupancy and Clustering Kinetics), a user selects and crops a nucleus of interest from SCAN’s output (cyan). Within that nucleus, LOCK tracks the transcription site across frames in both channels: the TF channel (top row), where the circled position marks the tracked locus, and the MCP channel (bottom row), where that same position is used to localize the transcription site itself. This lets LOCK test, frame by frame, whether a cluster overlaps the locus and whether the TF is enriched at the site. (D) Both ATAT modules write their measurements to CSV, alongside an analysis notebook that generates nucleus-wide cluster properties (e.g., average volume, enrichment, signal persistence), locus/gene-specific cluster properties (e.g., distance to site, enrichment, persistence), and representative images and movies for figures.

Three steps require user input. The first is recording each movie’s files, channels, and analyzed frames in a dataset table. The second is previewing and tuning cluster-size parameters on one representative frame before running SCAN on a full dataset. The third is choosing which nuclei to analyze in LOCK and manually reviewing and correcting locus tracks. All remaining steps can run automatically on a workstation or as batch jobs on a high performance computing (HPC) cluster using SLURM-scheduling. Dataset-specific settings (channels, frames, frame interval, voxel size, point-spread function (PSF)) are read from the dataset table and acquisition metadata. Settings shared across datasets (the enrichment threshold, and the displacement cutoffs that exclude fast-moving nuclei) are read from a per-machine configuration file.

ATAT writes its results as tables that can be plotted using a provided interactive notebook (Fig. 1D). SCAN produces one record per cluster and tracks volume, position, enrichment (cluster intensity relative to its own nucleus’s mean), and the half-life and zero-crossing time of the cluster’s intensity autocorrelation as measures of persistence. LOCK produces a record of TF enrichment at the locus over time, notes when the locus overlaps a segmented cluster, and calculates the persistence of enriched signal. Overall, these measurements can be used to assess relative occupancy at a site over time or the overall biophysical properties of TF clusters in a nucleus.

### Using SCAN to Segment and Quantify Nuclear TF Clusters

SCAN’s main function is to quantify TF clusters across the entire nuclear volume. Unlike global thresholding methods that are sensitive to fluctuations in protein concentrations, SCAN uses a local-contrast-based approach to identify and measure TF clusters. The first stage of SCAN involves defining the search space for cluster detection. Because the analysis is restricted to the nucleus, accurate nuclear segmentation is a prerequisite for normalizing cluster metrics. The framework uses 3D nuclear masks generated either through automated deep-learning-based segmentation using tools like CellPose^41^, or via a signal-intensity mask of a nuclear marker to isolate the nucleoplasm from the surroundings (Fig. 2A). Nuclei are then linked across frames using a nearest-neighbor algorithm. This masking step ensures that subsequent calculations of cluster density, defined as the number of clusters per unit volume, are accurately scaled to any changes in nuclear volume.

**Figure 2.**
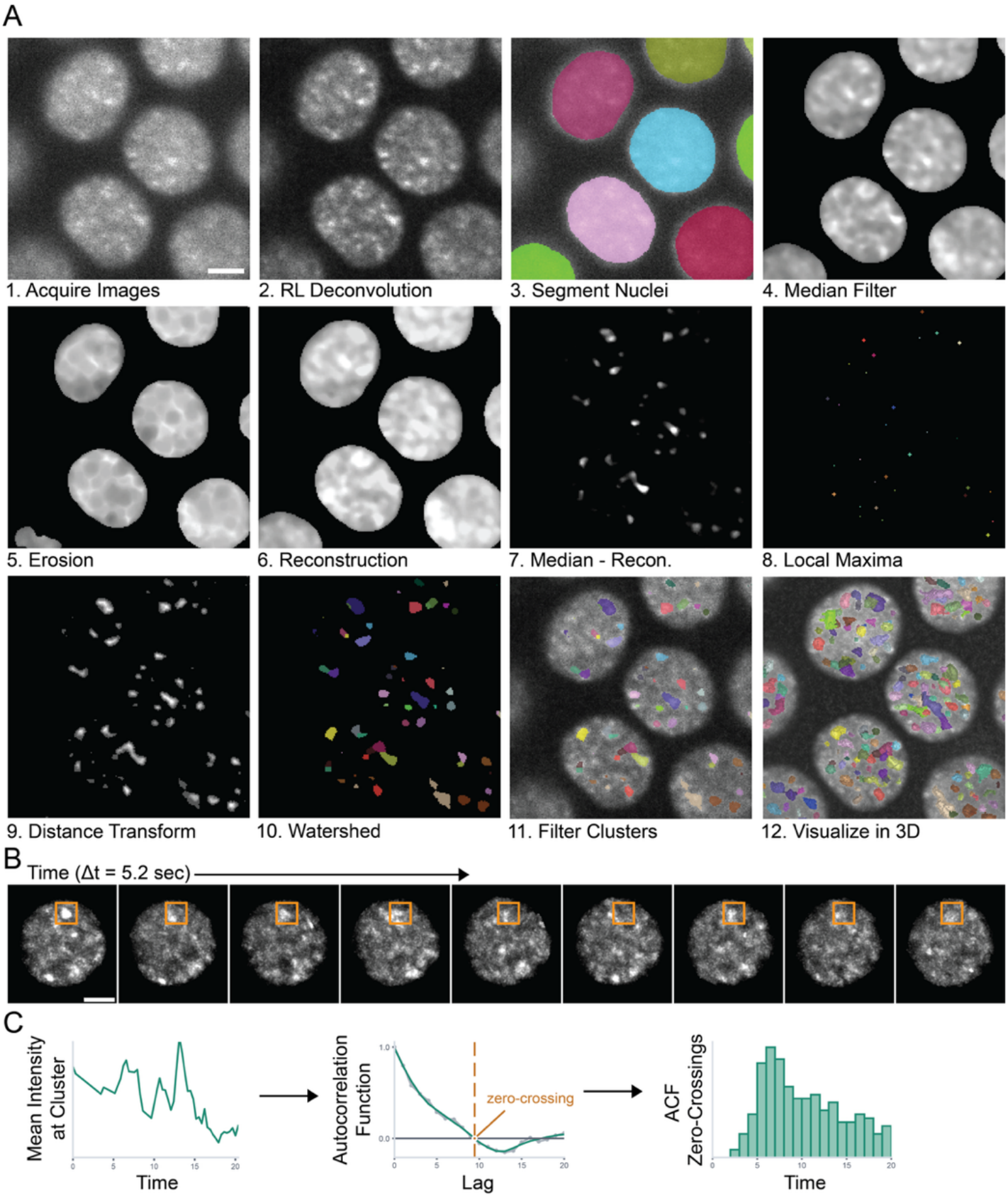
Automated pipeline for cluster segmentation and persistence using SCAN. (A) Stepwise visualization of the SCAN image analysis workflow. Initial input including (1) raw fluorescence image of mNeonGreen-Zelda in an nc13 *Drosophila* embryo, (2) Richardson-Lucy deconvolution, and (3) segmented and tracked nuclear labels. (4) Feature enhancement using a median filter followed by (5-7) morphological reconstruction-based background subtraction to isolate peaks (high-intensity candidates). (8) Identification of local maxima and (9) distance transform-seeded (10) watershed segmentation to resolve individual clusters. (11-12) Final cluster labels used for extraction of spatial coordinates, enrichment ratios, and more. Scale bar is 2 μm. (B) Representative time-lapse of a single nucleus (Zelda-mNeonGreen) over nine consecutive frames (Δt = 5.2 sec). Scale bar is 2 μm. To calculate cluster persistence, in a chosen frame (frame 1) a cluster’s centroid position is localized (orange box) and a window centered on that position is followed over subsequent frames. The mean intensity within this window is extracted to build a per-cluster intensity trace. (C) Quantification of cluster signal persistence begins with the mean intensity within the tracked window plotted over time. Then, the autocorrelation function (ACF) is calculated for this trace. The lag at which the smoothed ACF first crosses zero (dashed line) is taken as that cluster’s persistence time. The distribution of ACF zero-crossing times across all tracked clusters is plotted summarizing persistence behavior at the population level.

SCAN identifies clusters on the basis of local contrast and morphological features rather than absolute intensity. The workflow begins with the acquisition of raw fluorescence data, which then undergo GPU-accelerated Richardson-Lucy deconvolution^42^ to improve contrast and resolve near-diffraction-limited structures (Fig. 2A). These volumes are then integrated with tracked nuclear labels to constrain the analysis to the nucleoplasm, effectively excluding cytoplasmic noise and ensuring that each identified cluster is assigned to a specific parent nucleus.

SCAN uses a sequence of morphological operations to enhance features at the expected spatial scale of TF clusters (Fig. 2A). Following an initial median filter to suppress shot noise, the pipeline employs a morphological reconstruction-based background subtraction. This process involves eroding a filtered version of the image to create a marker, which is then iteratively dilated under the constraint of the filtered image (morphological reconstruction). By subtracting this reconstructed background from the original signal SCAN isolates local intensity peaks that serve as candidate clusters. To resolve individual clusters within dense environments, the software identifies local maxima within these peaks and applies a distance transform-seeded watershed segmentation. This approach enables cluster segmentation despite variations in overall nuclear intensity.

Once segmented, SCAN extracts a set of spatial and intensity measurements for each object, including 3D coordinates, volume, and enrichment ratios (defined as the mean intensity of the cluster normalized to the surrounding nucleoplasm). Beyond static morphology, the framework is equipped to quantify the temporal stability of these assemblies. To measure cluster persistence, SCAN defines a fixed window centered on a cluster’s position and extracts the mean intensity within that window over 20 subsequent frames (Fig. 2B). It then computes the autocorrelation function (ACF) of this intensity trace and smooths it to reduce frame-to-frame noise (Fig. 2C). From the smoothed ACF, SCAN reports the zero-crossing, the lag at which it first crosses zero. Zero-crossing times from all tracked clusters are then combined into a population-level distribution (Fig. 2C).

Because SCAN measures fluorescence within a fixed spatial window, its persistence metric reflects the temporal stability of local enrichment rather than the lifetime of an individually tracked cluster. A cluster moving out of the window can produce a loss of correlation even if it remains intact, while another cluster entering the same region may prolong the apparent persistence. To reduce the contribution of nuclear motion, nuclei exhibiting large displacements during the analysis window are excluded. These caveats are important to consider when interpreting absolute persistence measurements.

To validate the sensitivity and specificity of the SCAN segmentation pipeline, we compared the clustering profiles of a known transcription factor against a negative control for cluster formation and tested whether this distinction holds under increased imaging noise. We imaged mNeonGreen-Zelda, a pioneer factor known to form highly dynamic clusters in the early *Drosophila* embryo^8,43,44^, alongside a nuclear localization signal (NLS) fused to mNeonGreen, which serves as a benchmark for a spatially uniform, non-clustering protein distribution^32^. Each protein was imaged both at our standard excitation laser power (ZLD, NLS) and at approximately half laser power (ZLD^low^, NLS^low^) to generate a matched, noisier counterpart of each dataset. The NLS-tagged protein does not form discrete clusters under either imaging condition (Fig. 3A). While the initial cluster segmentation identifies candidate clusters in all four conditions, we applied an enrichment threshold (~1.3) calibrated against the NLS control so that its filtered cluster density approaches zero. This preserves Zelda clusters while returning a near-null result for both NLS conditions (Fig. 3A, D, F). Unfiltered cluster density and enrichment increase for both ZLD and NLS when imaging noise is increased (Fig. 3C, E), reflecting spurious noise that the initial segmentation alone cannot distinguish from true clusters, whereas after enrichment filtering this noise sensitivity is removed and the NLS result stays near zero regardless of imaging condition (Fig. 3D, F). Additionally, after filtering both Zelda imaging conditions give similar results. Taken together, the pipeline’s cluster segmentation is robust to differences in imaging quality.

**Figure 3.**
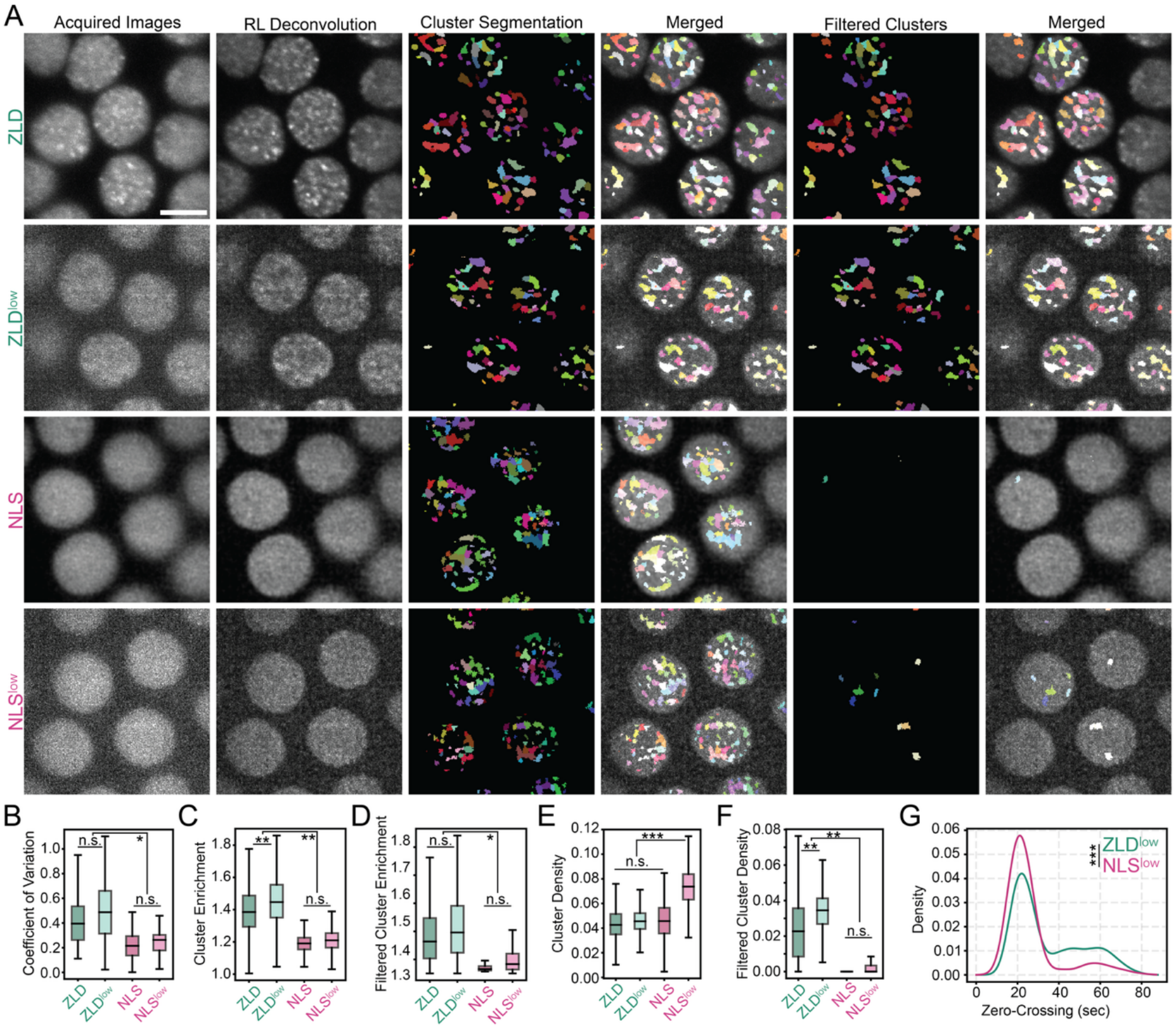
Validation of the SCAN pipeline using mNeonGreen-Zelda and NLS-mNeonGreen. (A) Representative max-intensity projections of *Drosophila* nc13 nuclei expressing mNeonGreen-Zelda (ZLD, top two rows) or NLS-mNeonGreen (NLS, bottom two rows), each imaged at normal laser power or at half power to increase image noise (ZLD^low^, NLS^low^). From left to right: raw acquired image, RL-deconvolved image, initial cluster segmentation, segmentation overlaid on the deconvolved image, segmentation after enrichment-threshold filtering, filtered segmentation overlay. Scale bar, 5 µm. (B-F) Quantification across all four conditions: (B) coefficient of variation of nuclear intensity, (C) mean enrichment of all detected clusters before filtering, (D) mean enrichment of clusters after enrichment-threshold filtering, (E) density of all detected clusters (clusters/µm^3^ nuclear volume), (F) density of clusters retained after filtering. (G) Distribution of cluster persistence (time to autocorrelation zero-crossing) for ZLD^low^ versus NLS^low^. Persistence was assessed only in the lower-laser-power acquisitions to avoid photobleaching confounds over time. Significance by two-sided Mann-Whitney U test (B-F) or two-sample Kolmogorov-Smirnov test (G): n.s., not significant; *p<0.05; **p<0.01; ***p<0.001. N = 16 ZLD, 11 ZLD^low^, 6 NLS, 9 NLS^low^ embryo replicates.

Quantitative analysis of these datasets further confirms that the SCAN detection engine is sensitive to higher-order protein organization rather than imaging noise alone. The intensity distribution of Zelda-expressing nuclei showed a significantly higher coefficient of variation (CV) than NLS-mNeonGreen (Fig. 3B), indicating a more heterogeneous, clustered signal. This CV difference held up between the noisier ZLD^low^ and NLS^low^ acquisitions while CV was not significantly different between the normal- and low-power acquisitions of either protein, indicating that the measure itself is robust to the increased imaging noise. After enrichment filtering, Zelda showed higher cluster density and cluster enrichment than NLS under both imaging conditions (Fig. 3D, F), consistent with the near-complete absence of filtered clusters in the NLS controls. Because persistence requires longer time-lapse acquisitions, we assessed it only in the lower-laser-power condition to avoid the risk of photobleaching confounding the result. Zelda clusters showed a significantly longer persistence than the sparse clusters detected in the NLS control (Fig. 3G). Together, these results demonstrate that SCAN can distinguish clusters from background fluctuations and provides reproducible measurements of nuclear clustering.

### Quantifying Locus-Specific TF Clusters with LOCK

While SCAN provides a statistical overview of the global nuclear landscape, understanding the formation and function of TF clustering requires measurements at individual genomic loci. LOCK quantifies the local accumulation of transcription factors at specific genomic targets by quantifying the average enrichment of a protein at that site relative to the nuclear background. This enrichment calculation is based on previous methods where the enrichment of a protein is averaged across timepoints or replicates^8,10,26,31,45–50^. Building on these methods, LOCK provides time-resolved analyses to study how the enrichment of a protein at a locus changes over time. The primary challenge in measuring enrichment is the need for high-precision spatial registration between the TF signal and a reporter for the gene of interest. LOCK addresses this by using a dual-channel analysis pipeline that centers its quantification on either nascent transcription sites marked by MS2/MCP fluorescent reporters or gene loci marked with other locus-labelling systems such as ParS/ParB or LacO/LacI arrays. By defining the gene locus as the coordinate origin, the framework can distinguish clusters that are preferentially enriched near the gene of interest.

The LOCK pipeline begins with the dual-channel acquisition of the transcription factor and in our example, the nascent transcripts of a target gene (Fig. 4A). After generating automated nuclear masks to exclude cytoplasmic signal, the software performs concurrent 3D segmentation of the MS2/MCP spot and comparison with the localization of clusters segmented using SCAN. To ensure accurate tracking of the labeled locus, LOCK uses a multi-stage workflow that requires user curation and input (Fig. 4B). Specifically, the detection phase begins with a maxima filter and difference-of-Gaussians (DoG) enhancement to identify candidate transcription sites, which are subsequently isolated using an adjustable percentile-based intensity threshold. To ensure the exclusion of non-specific spots and imaging artifacts, a filter is used to remove objects falling outside of biologically relevant size constraints. These validated sites are then linked across time frames using a nearest-neighbor tracking algorithm. If using a transcriptional reporter system such as MS2/MCP, the pipeline interpolates positions between active transcription bursts to maintain a continuous trajectory. Finally, manual tracking correction improves the accuracy of the genomic coordinates throughout the nuclear cycle. This approach allows for the persistent monitoring of a single locus even as it moves within the dynamic environment of the interphase nucleus (Fig. 4C).

**Figure 4.**
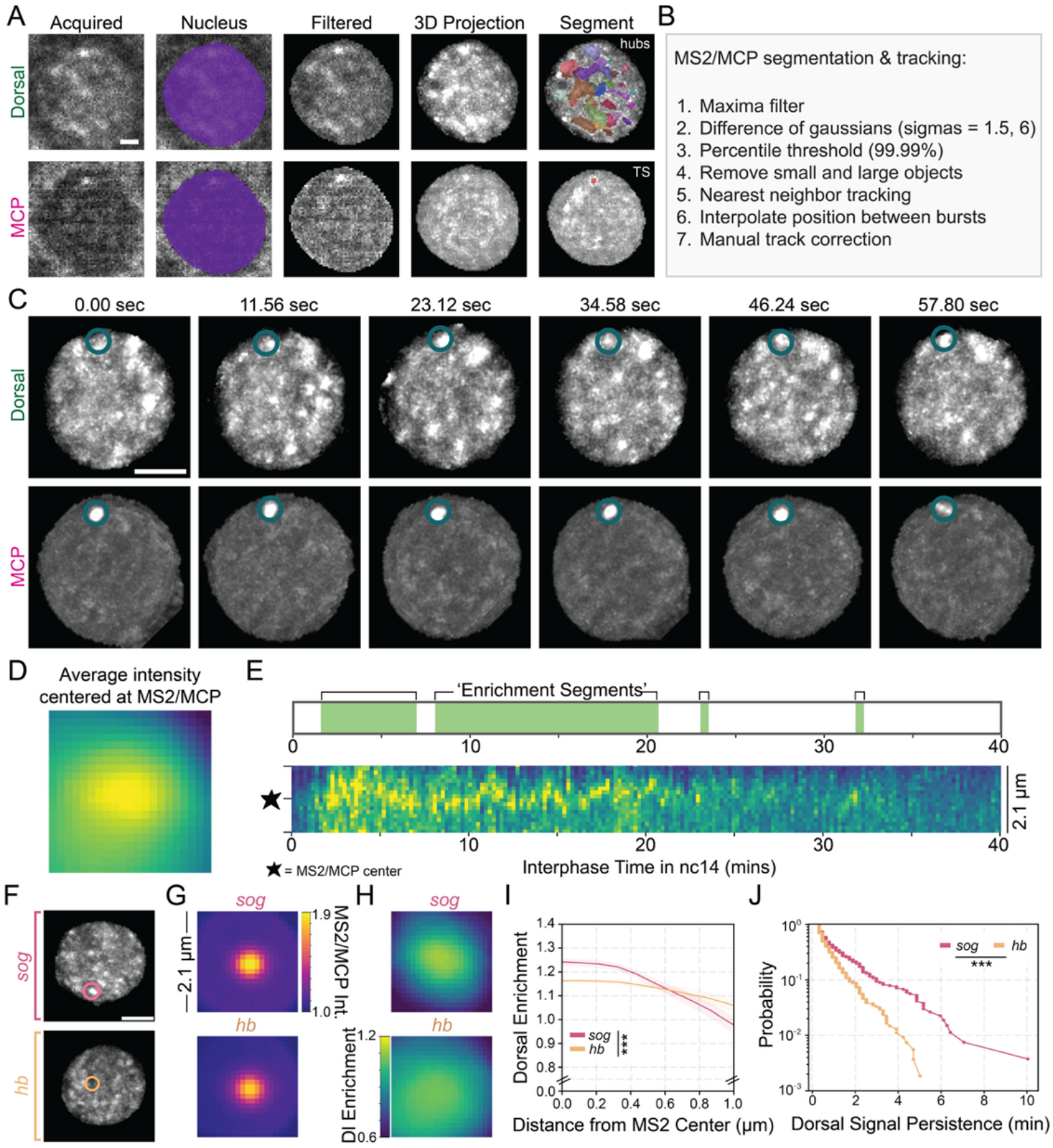
LOCK pipeline for tracking sites of nascent transcription and associated TF clusters. (A) Representative single-nucleus images of a *Drosophila* embryo in nc14 expressing Dorsal-mNeonGreen (top) and MCP-mCherry (bottom) to label nascent MS2 transcripts. From left to right: raw acquired fluorescence; automated nuclear mask generation; exclusion of cytoplasmic signal; 3D maximum intensity projection; and the resulting segmentation for Dorsal clusters and the MS2/MCP spot. Scale bar is 1 µm. (B) Schematic workflow detailing the pipeline for the segmentation and 4D tracking of MS2/MCP spot. (C) Time-lapse 3D projections of a single nucleus (Dorsal, top; MCP, bottom) over approximately 1 minute of interphase in nc14. Blue circles denote the persistent tracking of a single MS2/MCP spot across six consecutive frames. Scale bar is 2.5 µm. (D) Representative spatial average of Dorsal-mNeonGreen intensity centered on the coordinates of the tracked MS2/MCP spot, illustrating local enrichment. (E) (Bottom) Kymograph showing Dorsal-mNeonGreen intensity at the MS2/MCP locus throughout nc14 interphase. (Top) Schematic representation of “enrichment segments,” identifying discrete temporal windows where Dorsal intensity at the locus exceeds the nuclear mean. (F) Representative images of *Drosophila* nc14 nuclei expressing Dorsal-mNeonGreen. Circles denote the 3D coordinates of the tracked *sog* and *hb* MS2/MCP nascent transcription sites. Scale bar is 2 µm. (G) Regions centered on the MS2/MCP spot in the MCP-mCherry channel. (H) Corresponding cropped regions of the Dorsal-mNG channel at the same spatial coordinates as in (G). (I) Radial enrichment profiles extracted from the crops in (H), showing the mean normalized Dorsal-mNG intensity as a function of distance from the MS2/MCP spot center. Shaded regions represent the standard error of the mean. Center enrichment at the locus (0 µm) was compared between genes with a Mann-Whitney U test ***p<0.001. (J) Survival plots illustrating the distribution of “enrichment segment” lengths. Distributions of enrichment-segment lengths were compared with a two-sample Kolmogorov-Smirnov test on pooled segments ***p<0.001. N = 3 embryo replicates with 5 nuclei per embryo for both *sog* and *hb*.

Once the 3D trajectory of the transcription site is established, LOCK extracts the local fluorescence intensity of the transcription factor channel centered on those coordinates and also the overlap with SCAN-generated segmentation labels. To visualize the local TF fluorescence distribution, the software generates a spatial average of the TF intensity centered on the MS2/MCP spot, measuring local enrichment relative to mean nuclear fluorescence (Fig. 4 D). These spatial data are then extended into the temporal dimension to capture the dynamics of recruitment throughout the nuclear cycle. By generating a kymograph of the TF intensity at the tracked locus, we can observe how the local protein concentration fluctuates during interphase (Fig. 4E). To analyze these dynamics quantitatively, LOCK identifies “enrichment segments”, or discrete time blocks where the local TF intensity exceeds the nuclear mean in a persistent manner (Fig. 4E). This allows for the quantification of the persistence of TF enrichment at individual gene loci.

To validate LOCK, we tested its ability to resolve gene-specific TF clusters by comparing a known target gene against a non-target control. We used the TF Dorsal, endogenously tagged with mNeonGreen, and tracked the 3D coordinates of sites of nascent transcription of *short gastrulation (sog)*, a known Dorsal target gene, and *hunchback (hb)*, which serves as a negative control (Fig. 4F). By anchoring our analysis on the MS2/MCP nascent transcription spots (Fig. 4G), we could directly observe the local distribution of Dorsal-mNeonGreen in the immediate vicinity of these specific alleles (Fig. 4H). To quantify these spatial relationships, LOCK generates radial enrichment profiles that measure mean-normalized TF intensity as a function of distance from the center of the transcription site. The resulting profiles reveal a strong enrichment of Dorsal at *sog* that decays with increasing distance, confirming localized protein enrichment (Fig. 4I). In contrast, *hb* exhibits a markedly flatter profile near the nuclear mean intensity, confirming that this non-target locus lacks specific Dorsal enrichment. Beyond spatial enrichment, we used survival plots to analyze the duration of enrichment segments (Fig. 4J). The distribution of enrichment segment lengths shows that Dorsal clusters at *sog* display significantly greater temporal persistence than the transient fluctuations observed at the non-target *hb*. Together, these measurements demonstrate that LOCK resolves differences in the spatial enrichment and persistence of TFs at distinct genomic loci.

### Running ATAT

ATAT is distributed as a Python package with a graphical interface for every stage, built on napari^51^ for image display, alongside a complete command-line equivalent for each step. A single launcher script (atat.py) presents a menu for choosing a stage (preprocessing, SCAN, LOCK, or plotting) and then either opening that stage’s GUI, running one step on one sample, or submitting a batch of samples. Each stage’s GUI and its command-line scripts read the same underlying configuration, so either can be used interchangeably depending on the task and the machine available. Every step is driven by two files: a dataset CSV recording per-movie information (file paths, imaging channels, which nuclear-cycle frames to analyze, and which processing steps to run), and a single atat_config.yaml per machine generated interactively through each GUI’s Setup panel, recording whether that machine runs locally or submits to an HPC cluster via SLURM, where the data lives, and the microscope’s deconvolution parameters. The preprocessing GUI handles setup, dataset selection, and visual quality control of raw, deconvolved, and tracked-mask images, but launches deconvolution (which requires a CUDA-capable GPU) from the command line (or an HPC batch-submission script) rather than through the interface, while SCAN’s and LOCK’s GUIs run their steps directly since neither requires a GPU.

Three points in the pipeline are deliberately interactive rather than automated, and each has a dedicated GUI panel. In SCAN, a “Preview segmentation” panel lets a user load one frame and adjust the cluster-detection size parameters interactively before committing to a run across the full dataset. In LOCK, a “Pick nuclei” panel lets a user click to select which nuclei to analyze per nuclear cycle, and a dedicated MS2-tracking interface displays every detected locus as a clickable, color-coded track that can be merged, repositioned, or deleted to correct tracking errors before finalizing which track represents the true signal. Every other step, once these choices are made, can be run unattended, either locally or as a batch of HPC jobs, with one job submitted per sample. Full installation instructions, including for users with no prior coding experience, are provided in GETTING_STARTED.md; stage-specific usage is documented in each stage’s own README.

Once every sample has been run through preprocessing, SCAN, and LOCK, a single Jupyter notebook (Plotting_Manuscript_Figures.ipynb) turns their output into the manuscript figures, organized in the same order the pipeline itself runs: SCAN’s validation first, then LOCK’s. The SCAN section reads SCAN’s own output CSV and per-sample persistence files directly, groups every measurement by experimental condition, and produces nucleus-wide cluster enrichment over time, cluster density and cluster enrichment compared across conditions, and the ACF zero-crossing (persistence) distribution across conditions, each accompanied by a pairwise significance table. The LOCK section follows the same structure, grouped by gene, and produces mean-intensity crop images in both the TF and locus-reporter channels, radial enrichment profiles, enrichment at the locus over time, and an enrichment-segment survival curve.

Taken together, this design means a new user can run the complete pipeline without writing or modifying code. Every interactive decision happens through a GUI panel, every batch step reads the same configuration a GUI produces, and every figure and statistical comparison comes from a single notebook.

## DISCUSSION

ATAT was developed to provide consistent, automated measurements of nuclear protein clustering and its relationship to gene activity at specific genomic loci. By combining automated cluster segmentation with locus-specific measurements of protein enrichment, ATAT enables the spatial and temporal properties of nuclear clusters to be quantified across nuclei and at individual genes. Using Zelda and a nuclear-localized fluorescent protein as a control, we demonstrate that SCAN can distinguish TF clusters from background fluorescence fluctuations. We further show that LOCK resolves differences in Dorsal enrichment and persistence at distinct genomic loci.

Existing approaches to quantify transcription factor clustering often involve a trade-off between measurements across the entire nucleus and those at specific genomic loci. Segmentation-based analyses can characterize the spatial distribution, size, and enrichment of nuclear clusters, while locus-centered measurements can establish their relationship to specific genes. ATAT brings these approaches together by using SCAN to quantify nuclear clusters and LOCK to measure local enrichment and persistence at tracked genomic loci. Importantly, LOCK uses the same cluster segmentation generated by SCAN, allowing global and locus-specific measurements to be made using consistent criteria.

Although ATAT was developed using lattice light-sheet microscopy data from live *Drosophila* embryos, the underlying analysis is not specific to this imaging modality or model system. The combination of local-contrast-based cluster detection and locus-centered fluorescence measurements can be applied to other microscopy datasets in which nuclear protein distributions and genomic loci can be resolved. The integration of ATAT with napari provides an interactive interface for inspecting segmentation results and correcting locus trajectories, while command-line and batch-processing capabilities allow the analysis to be scaled across datasets without requiring users to modify the underlying code.

Our previous studies have used related approaches to connect nuclear protein clustering with transcription factor binding kinetics, enhancer architecture, and transcriptional activity^9,37,44,52^. ATAT consolidates these methods into a single, reusable toolkit that can be applied across proteins, genes, and experimental conditions. Importantly, the measurements provided by ATAT do not require assumptions about the mechanisms responsible for cluster formation. Rather, they provide quantitative descriptions of the spatial enrichment and temporal persistence of nuclear proteins that can be compared with models of molecular diffusion, chromatin binding, and higher-order protein interactions. By making these measurements more accessible and reproducible, ATAT provides a framework for investigating how nuclear protein organization relates to gene regulation.

### Limitations of the study

ATAT is limited by the spatial and temporal resolution and signal-to-background characteristics of the input microscopy data. In particular, the identification of clusters depends on the ability to distinguish local protein enrichment from fluorescence fluctuations arising from imaging noise. We therefore recommend including a non-clustering control, such as the nuclear-localized mNeonGreen used here, to establish an appropriate enrichment threshold. Because the optimal threshold depends on the properties of the fluorescent protein, its expression level, and the imaging conditions, it should be calibrated for each experimental system. ATAT was validated here using lattice light-sheet microscopy of live *Drosophila* embryos, and its performance with other imaging modalities and biological systems will depend on the quality and resolution of the available data.

An additional limitation concerns the interpretation of cluster persistence. SCAN calculates the autocorrelation of fluorescence intensity within a fixed spatial window rather than continuously tracking the identity of an individual cluster. Consequently, cluster motion can contribute to the loss of fluorescence correlation even when the underlying cluster remains intact. Conversely, different clusters occupying the same region at successive time points can contribute to sustained local enrichment. SCAN persistence measurements should therefore be interpreted as timescales of local fluorescence correlation rather than direct measurements of individual cluster lifetimes. LOCK addresses some of these limitations by anchoring fluorescence measurements to a tracked genomic locus, allowing local enrichment to be followed as the locus moves within the nucleus. However, LOCK requires an independently labeled genomic locus, and tracking may require manual correction, particularly when using transcriptional reporters that produce intermittent signals. Furthermore, neither approach can unambiguously distinguish individual clusters that approach, overlap, or separate at distances below the spatial resolution of the microscope. These limitations should be considered when relating fluorescence enrichment and persistence to the underlying molecular mechanisms of cluster formation.

## MATERIALS AND METHODS

### KEY RESOURCES TABLE

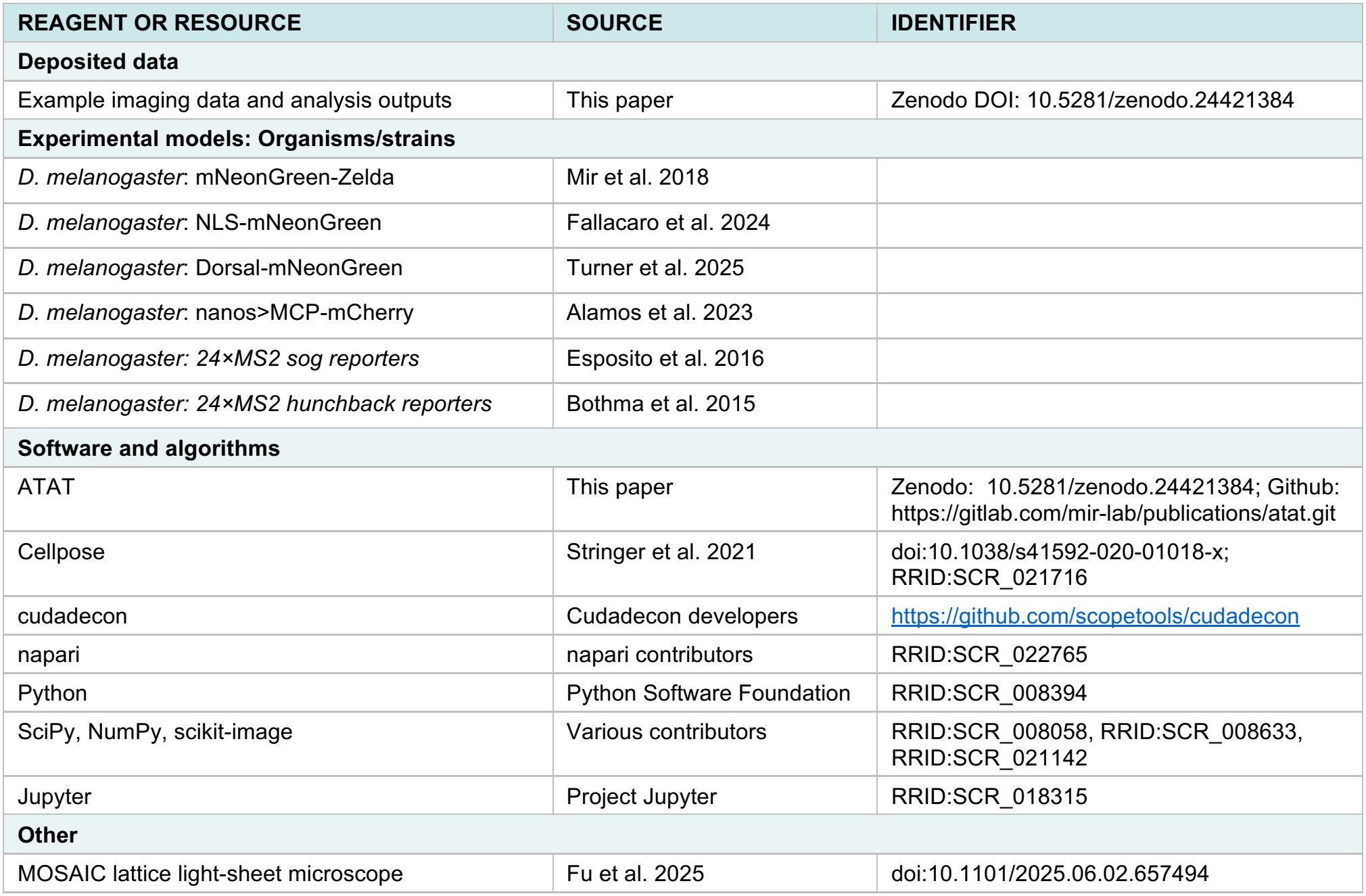

| **REAGENT OR RESOURCE** | **SOURCE** | **IDENTIFIER** |
|---|---|---|
| **Deposited data** |  |  |
| Example imaging data and analysis outputs | This paper | Zenodo DOI: 10.5281/zenodo.24421384 |
| **Experimental models: Organisms/strains** |  |  |
| *D. melanogaster*: mNeonGreen-Zelda | Mir et al. 2018 |  |
| *D. melanogaster*: NLS-mNeonGreen | Fallacaro et al. 2024 |  |
| *D. melanogaster*: Dorsal-mNeonGreen | Turner et al. 2025 |  |
| *D. melanogaster*: nanos>MCP-mCherry | Alamos et al. 2023 |  |
| *D. melanogaster: 24×MS2 sog reporters* | Esposito et al. 2016 |  |
| *D. melanogaster: 24×MS2 hunchback reporters* | Bothma et al. 2015 |  |
| **Software and algorithms** |  |  |
| ATAT | This paper | Zenodo: 10.5281/zenodo.24421384; Github: https://gitlab.com/mir-lab/publications/atat.git |
| Cellpose | Stringer et al. 2021 | doi:10.1038/s41592-020-01018-x; RRID:SCR_021716 |
| cudadecon | Cudadecon developers | https://github.com/scopetools/cudadecon |
| napari | napari contributors | RRID:SCR_022765 |
| Python | Python Software Foundation | RRID:SCR_008394 |
| SciPy, NumPy, scikit-image | Various contributors | RRID:SCR_008058, RRID:SCR_008633, RRID:SCR_021142 |
| Jupyter | Project Jupyter | RRID:SCR_018315 |
| **Other** |  |  |
| MOSAIC lattice light-sheet microscope | Fu et al. 2025 | doi:10.1101/2025.06.02.657494 |

### EXPERIMENTAL MODEL AND STUDY PARTICIPANT DETAILS

#### Drosophila melanogaster

To visualize transcription factor (TF) dynamics and nascent transcription, several endogenous and transgenic *Drosophila melanogaster* lines were used. The pioneer factor Zelda was visualized using an endogenously tagged mNeonGreen-Zelda line (mNG-ZLD) previously generated^8^, and a nuclear-localized monomeric mNeonGreen (NLS-mNG) we generated previously^44^ served as a spatially uniform control for background noise and stochastic fluctuation. For experiments involving the MS2/MCP system, virgin females expressing the tagged TF (Dorsal-mNeonGreen previously characterized^9,53^) and MCP-mCherry (expressed under the *nanos* promoter) were crossed to males carrying MS2 reporters. These reporters contained 24 copies of the MS2 stem-loop sequence positioned downstream of *sog* or *hunchback* enhancers (MCP and reporter lines from previous work^54^). All flies were maintained at 25°C. Embryos were collected on apple juice agar plates supplemented with yeast paste. After a 1.5-2 hour collection period, embryos were dechorionated in 50% bleach for 2 minutes, rinsed thoroughly with deionized water, and mounted onto coverslips for lattice light-sheet Microscopy.

### METHOD DETAILS

#### Lattice Light-Sheet Imaging

All imaging was performed on a MOSAIC system^55^ in the lattice light-sheet mode, as previously described, optimized for fly embryo imaging^37^. Volumetric imaging was performed during nuclear cycles 13 (nc13) and 14 (nc14). In the SCAN validation (Figs. 2, 3), data were acquired in the higher 488 nm laser power setting of ~0.55 W and in the lower 488 nm laser power setting of ~0.25 W at a temporal resolution of ~5.2 seconds per volume. In the LOCK examples (Fig. 4), data were acquired with ~0.15 W of the 488nm laser and ~0.27 W of a 589 nm laser at a temporal resolution of ~11.56 seconds per volume. To minimize phototoxicity while maintaining high signal-to-noise ratios, exposure times were limited to 30 ms per slice with a 0.3 µm Z-step.

#### Image Preprocessing

Prior to cluster detection, all raw volumetric data underwent GPU-accelerated Richardson-Lucy deconvolution (5 iterations)^42^ using a measured point-spread function (PSF). Every nucleus in each deconvolved volume was segmented using a pretrained Cellpose model and linked across frames by a nearest-neighbor tracking algorithm (maximum per-frame displacement of 50 pixels, no permitted frame gap), producing a tracked nuclear mask for each embryo that both SCAN and LOCK read directly rather than re-deriving. Users may substitute deconvolved images and nuclear label matrices generated by an alternative segmentation method in place of this step, provided they are supplied in the same per-frame, per-nucleus-labeled format.

#### ATAT: Software Implementation and Usage

ATAT is implemented in Python and organized as three sequential stages (preprocessing, SCAN, LOCK) plus a plotting notebook, each with a graphical interface built on napari for interactive steps and a complete command-line equivalent for batch execution, including submission to an HPC cluster via SLURM. Every stage reads from a shared, per-dataset CSV recording each movie’s file paths, imaging channels, analyzed frames, and condition/genotype labels, and a per-machine configuration file recording acquisition parameters and run-mode (local vs. HPC) settings. This separation allows the same analysis code to run unmodified across machines and datasets without requiring the user to edit source code. Software and sample-data availability are given under Resource Availability. The code repository includes README files with detailed step-by-step instructions.

#### SCAN: Segmentation and Clustering Analysis in Nuclei

The SCAN software package was developed in Python to automate the 3D detection and characterization of TF clusters within already-segmented, tracked nuclei. For every frame, each nucleus’s TF signal is first normalized to that nucleus’s own mean intensity, so that clusters are called relative to each nucleus’s own signal rather than against a single intensity cutoff applied across the dataset. The pipeline then applies a median filter to suppress shot noise, followed by a morphological reconstruction-based background subtraction (erosion of the filtered image to generate a marker, followed by grayscale reconstruction by dilation of that marker under the filtered image) to isolate high-intensity peaks from the normalized, locally varying nucleoplasmic background. Local maxima within these peaks seed a distance transform-based watershed segmentation to resolve individual clusters. Candidate objects below a minimum voxel size are discarded. For each segmented object, SCAN calculates volume, 3D centroid coordinates, and enrichment ratios (mean cluster intensity relative to the surrounding nucleoplasm, computed for raw, deconvolved, and normalized intensity independently). Cluster density is calculated as the number of clusters per unit nuclear volume, with nuclear volume taken from the same tracked nuclear mask used for normalization.

A single enrichment threshold is applied to this candidate cluster population, once per dataset, at nuclear cycle 14, to distinguish true clusters from residual background fluctuations surviving the morphological segmentation. This threshold is calibrated empirically for each protein, fluorophore, expression regime, and imaging system by comparing the filtered-cluster yield in the experimental condition against a non-clustering control (here, NLS-mNeonGreen). The threshold is chosen such that it reduces the non-clustering control’s filtered cluster count, density, and enrichment to near zero while preserving detection of clusters in the clustering condition and is then held fixed for all comparisons within that dataset. Reporting both the unfiltered and filtered populations makes this specificity directly visible. The unfiltered population reflects every locally bright region the morphological segmentation step can resolve, including noise, whereas the filtered population isolates only those regions enriched above what the non-clustering control produces at the same threshold.

Persistence of each filtered cluster is quantified by extracting the mean intensity within a 1 µm window centered on the cluster’s position over 20-frames, computing the autocorrelation function (ACF) of that intensity trace, and smoothing the result with a Savitzky-Golay filter. Both the half-life (the lag at which the smoothed ACF falls to half its initial value) and the zero-crossing (the lag at which it first crosses zero) are reported from the smoothed ACF by linear interpolation. The zero-crossing is the persistence metric reported throughout the Results. Nuclei that move more than 0.6 µm axially or 0.3 µm laterally between frames are excluded from this calculation to avoid attributing an apparent loss of correlation caused by nuclear motion to a change in signal.

Per-nucleus skewness, kurtosis, and coefficient of variation of the normalized intensity distribution, computed across all voxels within each nuclear mask were calculated as an independent, segmentation-free measure of intensity heterogeneity, used alongside the segmentation-based cluster metrics above to validate detection specificity.

#### LOCK: Locus-specific Occupancy and Clustering Kinetic

The LOCK package was developed to measure the local protein microenvironment at specific genomic loci, anchored on a transcription site identified in a second imaging channel. Candidate transcription-site spots are detected with a difference-of-Gaussians filter (default σ = 1.5 and 6.0 pixels but adjustable in the GUI) after a maxima filter and nuclear-mask-based exclusion of cytoplasmic signal, followed by a percentile-based intensity threshold (default 99.9th percentile, also adjustable in the GUI) and removal of objects below a minimum size. Detected spots are linked frame-to-frame using the same nearest-neighbor tracking algorithm used for nuclear tracking during preprocessing. Resulting tracks are then manually reviewed and corrected as needed.

Once a locus trajectory is established, LOCK extracts the local fluorescence intensity of the TF channel at that position and tests, frame by frame, whether the locus position overlaps a cluster mask produced by SCAN for that same nucleus (rather than re-segmenting clusters independently), and extracts a 3D radial fluorescence profile centered on the locus to quantify TF enrichment as a function of distance. An “enrichment segment” is defined as a run of consecutive frames in which TF intensity at the locus exceeds the nuclear mean and the locus position overlaps a SCAN-defined cluster mask by at least one voxel, allowing a single-frame gap before closing out a segment; the distribution of segment durations is reported as a survival curve.

### QUANTIFICATION AND STATISTICAL ANALYSIS

For comparisons between conditions (Fig. 3B–F) or genes (Fig. 4), each metric was averaged to a single value per embryo. Significance was assessed with a two-sided Mann-Whitney U test on the embryo-level means, with Bonferroni correction across all tests within a comparison. Mann-Whitney was chosen because it doesn’t assume normality. For persistence (zero-crossing) distributions (Fig. 3G) and enrichment-segment durations (Fig. 4J), a two-sample Kolmogorov-Smirnov test was run on pooled per-cluster values and reported as a supplementary, non-embryo-corrected check. Significance is denoted *p < 0.05, **p < 0.01, ***p < 0.001. Exact n, the definition of n, the center (median or mean) and the dispersion (SD, SEM, or interquartile range) are given in the figure legends. All tests were run with scipy.stats in Plotting_Manuscript_Figures.ipynb available in the provided code.

## ADDITIONAL RESOURCES

- ATAT installation guide (GETTING_STARTED.md) and stage-specific documentation: https://gitlab.com/mir-lab/publications/atat.git.
- Example dataset to reproduce Figs 3 and 4 doi:10.5281/zenodo.24421384

## ACKNOWLEDGMENTS

We thank all members of the Mir Lab for insightful discussion and feedback as testers of these packages on a variety of nuclear protein clusters.

## AUTHOR CONTRIBUTIONS

Conceptualization, S.F. and M.M.; methodology, S.F. and M.M.; investigation, S.F. and M.M.; writing – original draft, S.F. and M.M.; writing – review & editing, S.F. and M.M.; funding acquisition, S.F. and M.M.; supervision, M.M.

## DECLARATION OF GENERATIVE AI AND AI-ASSISTED TECHNOLOGIES in the writing process

The authors declare the use of generative AI in the research and writing process. According to the GAIDeT taxonomy (2026), the following tasks were delegated to GAI tools under full human supervision: code optimization, process automation, creation of algorithms for data analysis, visualization, proofreading and editing, and reformatting. The GAI tool used was: Claude Sonnet 5.5. Responsibility for the final manuscript lies entirely with the authors. GAI tools are not listed as authors and do not bear responsibility for the final outcomes. Declaration submitted by: Samantha Fallacaro

## RESOURCE AVAILABILITY

### Lead contact

Requests for further information and resources should be directed to and will be fulfilled by the lead contact, Mustafa Mir.

### Materials availability

This study did not generate new unique reagents.

### Data and code availability

- All data reported in this paper will be shared by the lead contact upon request.
- All original code has been deposited at Zenodo and is publicly available at doi: 10.5281/zenodo.24421384 as of the date of publication. Code can also be accessed through Gitlab: https://gitlab.com/mir-lab/publications/atat.git
- Any additional information required to reanalyze the data reported in this paper is available from the lead contact upon request.

